# Diversity of funnel plasmodesmata in angiosperms: the impact of geometry on plasmodesmal resistance

**DOI:** 10.1101/2021.05.05.442713

**Authors:** Grayson P. Ostermeyer, Kaare H. Jensen, Aslak R. Franzen, Winfried S. Peters, Michael Knoblauch

## Abstract

In most plant tissues, threads of cytoplasm, or plasmodesmata, connect the protoplasts via pores in the cell walls. This enables symplasmic transport, for instance in phloem loading, transport, and unloading. Importantly, the geometry of the wall pore limits the size of the particles that may be transported, and also (co-)defines plasmodesmal resistance to diffusion and convective flow. However, quantitative information on transport through plasmodesmata in non-cylindrical cell wall pores is scarce. We have found conical, funnel-shaped cell wall pores in the phloem-unloading zone in growing root tips of five eudicot and two monocot species, specifically between protophloem sieve elements and phloem pole pericycle cells. 3D reconstructions by electron tomography suggested that funnel plasmodesmata possess a desmotubule but lack tethers to fix it in a central position. Model calculations showed that both diffusive and hydraulic resistance decrease drastically in conical cell wall pores compared to cylindrical channels, even at very small opening angles. Notably, the effect on hydraulic resistance was relatively larger. We conclude that funnel plasmodesmata generally are present in specific cell-cell interfaces in angiosperm roots, where they appear to facilitate symplasmic phloem unloading. Interestingly, cytosolic sleeves of most plasmodesmata reported in the literature do not resemble straight annuli but possess variously shaped widenings. Our evaluations suggest that widenings too small for identification on electron micrographs may drastically reduce the hydraulic and diffusional resistance of these pores. Consequently, theoretical models assuming cylindrical symmetries will underestimate plasmodesmal conductivities.

## Introduction

Cells in plant tissues are defined by cell walls, rigid extracellular networks consisting of polysaccharides and, to a lesser degree, proteins. The cells are not physically isolated, though, as they are connected by thin cytoplasmic threads that extend through pores in the walls. The size and structure of these plasmodesmata can be modified by the living cells depending on their physiological and developmental requirements (Peters et al. 2021). Plasmodesmata may be simple (i.e., cylindrical), branched, or of more complex, often asymmetric geometries (Lee and Frank 2018). The interfaces between certain cell types may be characterized by specific plasmodesmal structures. For instance, the walls forming the interface between the sieve elements and companion cells in the phloem are perforated by so-called pore-plasmodesma units, branching cell wall pores with a large opening on the side of the sieve element and several smaller openings facing the companion cell (Esau and Thorsch 1985).

Plasmodesmata enable diffusion and in some cases bulk flow between cells. Such movement can be visualized using fluorescent reporter techniques including fluorochrome microinjection (Goodwin et al. 1990; Barton et al. 2011), fluorescence recovery after photobleaching (Wang et al. 2020; Rutschow et al. 2011), photoactivatable fluorochromes (Liesche and Schulz 2012), and photo-inducible fluorescent proteins (Gerlitz et al. 2018). Obviously, the size of particles that may move through a plasmodesma is limited by the dimensions of the plasmodesma; plasmodesmata function as sieves, as it were (Schulz 1999). Size exclusion limits below 1 kDa have been reported from tissues in photoassimilate-exporting leaves that often possess branched plasmodesmata (Oparka et al. 1999). Simple plasmodesmata seem to be involved in a specific assimilate export mechanism known as polymer trap, which requires size exclusion limits around 0.5 kDa (Comtet et al. 2017). In contrast, larger molecules of up to 40 kDa appear to move through simple plasmodesmata in photoassimilate-importing tissues (Oparka et al. 1999; Nicolas et al. 2017; Lee and Frank 2018), while pore-plasmodesma units enable the exchange of probes of up to 70 kDa (Oparka and Turgeon 1999; Fitzgibbon et al. 2013). Funnel-shaped plasmodesmata in the root unloading zone of *Arabidopsis* permit movements of molecules of at least 112 kDa (Ross-Elliott et al. 2017).

The developmental and cell type-specific variation of plasmodesmal size exclusion limits indicates active regulation and thus physiological significance of the effective size of the plasmodesmal pore. Therefore the geometry of plasmodesmata must be expected to play a role in controling cell-to-cell conductivity. Most theoretical plasmodesma models for quantitative evaluations generally assumed coaxial symmetry with a straight, cylindrical cell wall pore, based on interpretations of electron micrographs by e.g. Ding et al. (1992) and Waigmann et al. (1997). Consequently, models including a cytosolic sleeve mostly assumed this sleeve to be tubular with constant radius and annular cross-sectional shape, or to consist of a group of circularly arranged cylindrical tubes (Comtet et al. 2017; Liesche and Schulz 2013; Park et al. 2019). In reality, however, asymmetric and irregular shapes are common. For example, central cavities – diameter widenings in the center of plasmodesmata – have been described repeatedly (Ding et al. 1992; Nicolas et al. 2017; Fitzgibbon et al. 2010), but rarely were considered in theoretical analyses of plasmodesmal transport. Blake (1978) modeled convective flow but not diffusion in plasmodesmata that widened in the center, while Deinum et al. (2019) analyzed diffusion but not convective flow. Electrical effects have never been evaluated, although increasing plasmodesmal diameter at constant Debye length will reduce the influence of static wall charges on the movement of charged particles in the cytosol (Peters et al. 2021).

Root growth is fueled by materials that are unloaded from protophloem sieve elements (PSEs), the youngest fully functional components of the sieve tubes that reach the root elongation zone. There are two such tubes in *Arabidopsis*, and the unloading sieve elements typically have five neighbors: two companion cells, an immature metaphloem sieve element, and two phloem pole pericycle cells (PPPs). The PSE/PPP interfaces are characterized by unusual, funnel-shaped plasmodesmata (Ross-Elliott et al. 2017); apparently analogous plasmodesmata in *Hordeum vulgare* had been briefly discussed by Warmbrodt (1985). Initial calculations indicated that the conical funnel shape facilitated convective phloem unloading at low pressure differentials (Ross-Elliott et al. 2017). These findings supported the idea that the PSE/PPP interface has a specific significance for phloem unloading in *Arabidopsis* roots.

Since phloem unloading mechanisms are of major importance in the context of food production for human and lifestock consumption, detailed knowledge of the distribution of funnel plasmodesmata in species other than *Arabidopsis* as well as a better understanding of the physics of transport through non-cylindrical plasmodesmata would seem desirable. Therefore, and because funnel plasmodesmata appeared a convenient case for studying the impact of non-cylindrical pore shapes on plasmodesmal transport, we first established the general occurrence of funnel plasmodesmata in angiosperms. Then, we generated 3D reconstructions based on electron tomograms of these plasmodesmata to evaluate flow patterns and resistances, and modeled physical flow characteristics in idealized cylindrical and conical plasmodesmata, to evaluate the effects of various pore geometries. As a general conclusion, we suggest that often overlooked deviations from a straight, cylindrical shape can reduce the plasmodesmal resistance to bulk flow and diffusion significantly.

## Results

### Funnel plasmodesmata in seven species: basic observations

To investigate the ultrastructure of the interface of the protophloem with surrounding cells, we adapted microwave-supported fixation protocols for five eudicots from various families and two Poaceae species as representatives of the monocots. Variations in root thickness, tissue density, and other structural parameters made it necessary to adjust protocols for several species (see Methods). We note that protophloem sieve elements in roots of most species are more difficult to preserve for electron microscopy than those in the thin roots of *Arabidopsis*. Vitrification by cryo-fixation usually fails as the protophloem is located too deep within the organ to achieve the required freezing speeds.

With few exceptions, eudicots exhibit a bi-, tri-, tetra-, or pentarch architecture of the primary root, i.e. the roots possess two, three, four, or five protoxylem/protophloem units. In contrast, Poaceae roots generally are polyarch, showing a greater number of protophloem strands. Our study species confirmed this pattern (Fig. 1A, tetrarch root of *Ipomoea nil*; Fig. 1B, polyarch root of *Triticum aestivum*). Viewed on cross-sections of the zone of phloem unloading, the protophloem sieve elements (PSEs) of the eudicots examined were connected to five cells (Fig. 1C). These comprised two phloem pole pericycle cells (PPPs), two companion cells (CCs), and one immature metaphloem sieve element (MP). In contrast, protophloem sieve elements had only four direct neighbors in the Poaceae examined, as they were not reached by metaphloem sieve elements in the phloem unloading zone (Fig. 1D).

**Figure 1:**
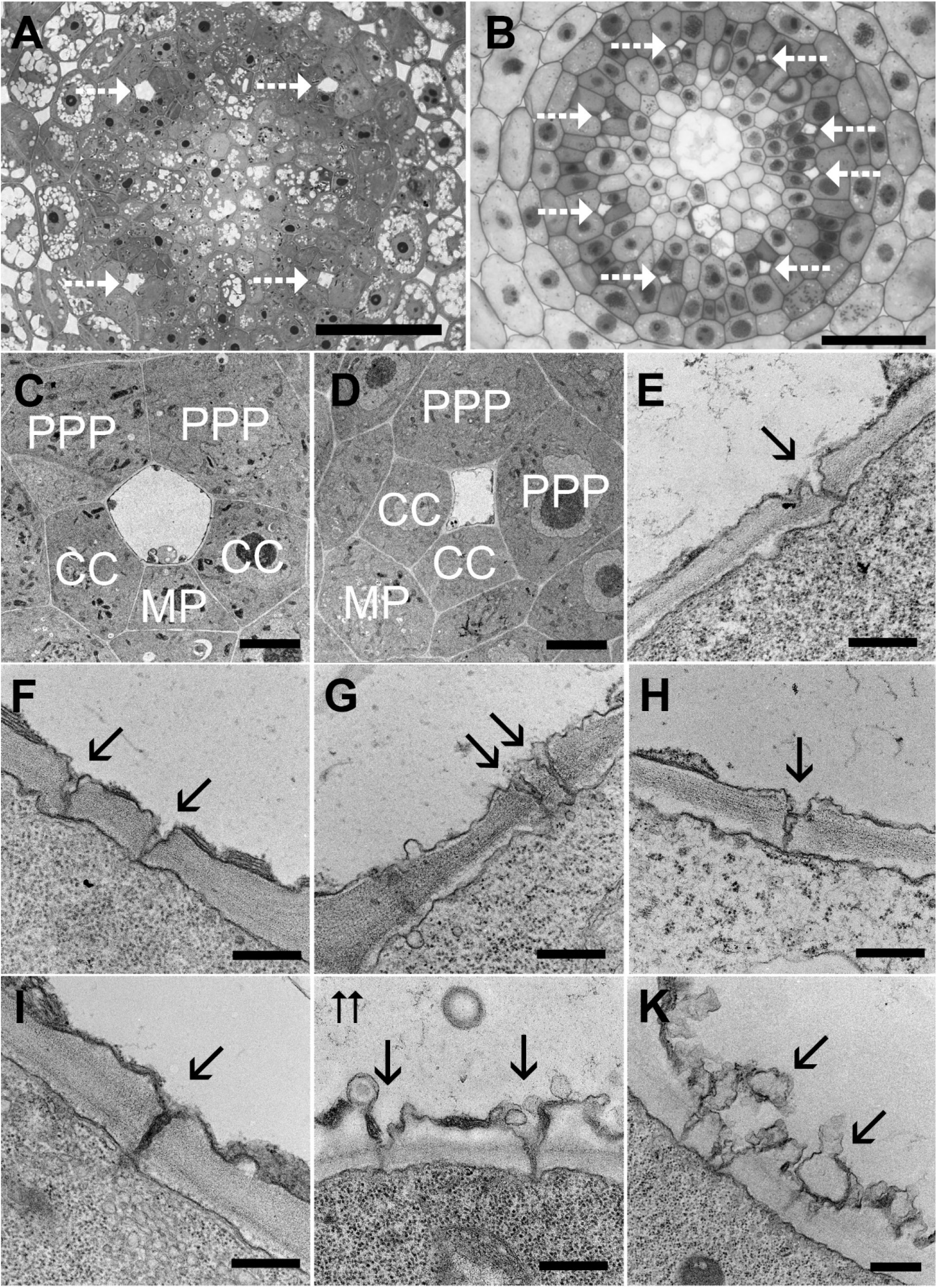
Tissue structure and funnel plasmodesmata in phloem unloading zones of growing root tips. A) Cross-section of the central cylinder in the phloem unloading zone in a root of *Ipomoea nil*, representative of the eudicots examined. The tetrarch vascular system has four protophloem sieve elements, which appear empty due to the absence of dense cytoplasm (dashed arrows). B) Analogous section of a polyarch root of *Triticum aestivumhere* with eight protophloem sieve elements (dashed arrows). C) Protophloem sieve elements typically connect to two companion cells (CC), two phloem pole pericycle cells (PPP), and one immature metaphloem sieve element (MP) in eudicots, shown here in *Ipomea nil*. D) In phloem unloading zones of the Poaceae roots tested, immature metaphloem sieve elements do not reach the protophloem sieve elements, leaving them with only four direct neighbors. The example shown is from *Oryza sativa*. In both (C) and (D), straight cell walls and the coherent structure of the cytoplasm indicate excellent preservation. E-K) Funnel plasmodesmata (solid arrows) in the PSE/PPP interfaces of five eudicots (E, *Ipomoea nil*; F, *Nicotiana tabacum*; G, *Catalpa speciosa*; H, *Solenostemon scutellarioides*; I, *Medicago sativa*) and two Poaceae (J, *Oryza sativa*; K, *Triticum aestivum*). Scale bars: A,B: 100 μm; C,D: 5 μm; E-H: 400 nm; K: 500 nm.

We found funnel-shaped plasmodesmata in all species examined (Fig. 1E–K). As in *Arabidopsis*, the occurrence of these funnel plasmodesmata was restricted to the walls between protophloem sieve elements and phloem pole pericycle cells, the PSE/PPP interfaces. The surfaces of these walls appeared relatively smooth in the eudicots (Fig 1E-I). In the Poaceae, the wall surfaces on the side of the sieve elements were rough with numerous protrusions and invaginations (Fig. 1J, K). In all species, the wider apertures of funnel plasmodesmata were always on the side of the protophloem sieve element (Fig. 1E–K), and thus served as inlet apertures of the plasmodesmata in phloem unloading.

### Geometry of funnel-shaped cell wall pores

In general, the geometries of the funnel-shaped cell wall pores were somewhat irregular due to their rugged inner surfaces. Nonetheless, the diameter, *D*_PSE_, of the wider plasmodesmal aperture on the side of the protophloem sieve element as well as the length, *L*, of the plasmodesma (equivalent to cell wall thickness) could be determined on numerous micrographs (*n* = 18 for *S. scutellarioides, n* = 30 for all other species; examples are shown in Fig. 1E-K, and data are compiled in Supplemental Table S1). If the opening angles, θ, of the funnel-shaped cell wall pores were constant, *D*_PSE_ should increase with increasing *L* for geometrical reasons (Fig. 2A). However, there was no correlation between the measured values of *D*_PSE_ and *L* (Fig. 2B), suggesting that θ should decrease with *L*. The estimation of θ was difficult as the narrow aperture diameters facing the phloem pole pericycle cells, *D*_PPP_, could not be determined unequivocally on many of the micrographs. Because our rough estimates of *D*_PPP_ on all analyzed micrographs ranged between 17 and 51 nm, we calculated θ for each plasmodesma with *D*_PPP_ set to 20 and 50 nm, to cover the range of realisticly expectable values. In fact, θ showed a tendency to decline with increasing wall thickness as indicated by the Geometric Mean Functional Relationship (GMFR) between the two parameters (Fig. 2C), but this relationship was far too weak to support general conclusions (*r*^2^ = 0.12 for data calculated with *D*_PPP_ = 20 nm, and *r*^2^ = 0.08 for *D*_PPP_ = 50 nm). While θ varied widely, there were species-specific trends; the median of θ determined separately for each species was smallest in *C. speciosa* and five to ten times higher in *O. sativa*, dependening on which value of *D*_PPP_ was assumed (Fig. 2D).

**Figure 2:**
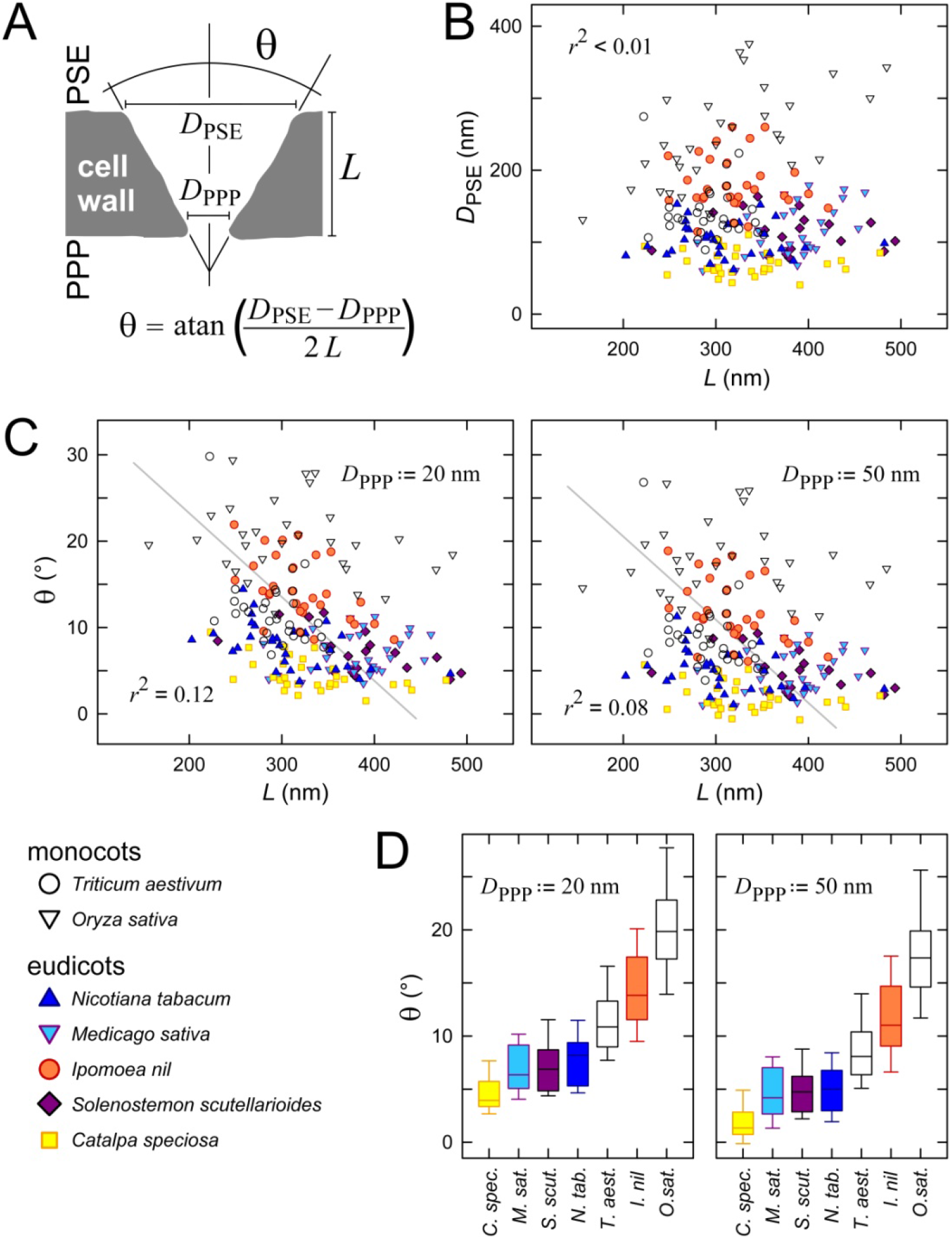
Cell wall pore geometry in the PSE/PPP interfaces of different species. (A) Pore opening angles (θ) were computed based on measurements taken on electron micrographs of plasmodesma length (equivalent to cell wall thickness), *L*, and the aperture diameter on the side of the protophloem sieve element, *D*_PSE_. The diameter of the aperture facing the phloem pole pericycle, *D*_PPP_, was set to 20 or 50 nm. (B) No correlation existed between measurements of *D*_PSE_ and *L* (*r*^2^ < 0.01). (C) Relation between *L* and pore opening angle θ, assuming a *D*_PPP_ of 20 nm (left) and 50 nm (right). Grey lines indicate the Geometric Mean Functional Relationship (GMFR). (D) Distribution of pore opening angles θ in different species (*D*_PPP_ set to 20 nm and 50 nm as indicated). Boxes represent the second and third quartile of the data with the median given as a horizontal line; whiskers indicate the 10^th^ and 90^th^ percentiles. The number of plasmodesmata analyzed was *n* = 18 for *S. scutellarioides* and *n* = 30 for all other species (see Supplemental Table S1 for original data).

### 3D reconstructions of individual plasmodesmata

Our above considerations of the geometries of funnel-shaped cell wall pores were based on the simplifying assumptions presented in Fig. 2A. To obtain a more realistic picture of the structure of individual funnel plasmodesmata, we generated 3D reconstructions by electron tomography. We acquired 220 nm slices of stained, resin-embedded samples for all species except *O. sativa*, which required 280 nm sections, wide enough to carry complete funnel plasmodesmata. Our reconstructions showed funnel plasmodesmata of varying, irregular shapes (Fig. 3). While we were able to identify desmotubules, it must be cautioned that the apparent diameters of these structures depend on parameters including sample thickness, resin hardness, desmotubule location, staining time, etc. In none of our image series, desmotubules appeared as sharply bordered structures but rather showed as gradients of contrast intensity. Therefore their exact dimensions were not always clearly discernible, and variations in apparent desmotubule diameter might reflect methodological uncertainty as much as natural variability of the actual structure. Given this caveat, it still appears noteworthy that funnel plasmodesmata showed no signs of tether-like connections between desmotubule and plasma membrane. Rather than being fixed in the center of the pore, the location of desmotubules in our reconstructions was highly variable (Fig. 3).

**Figure 3:**
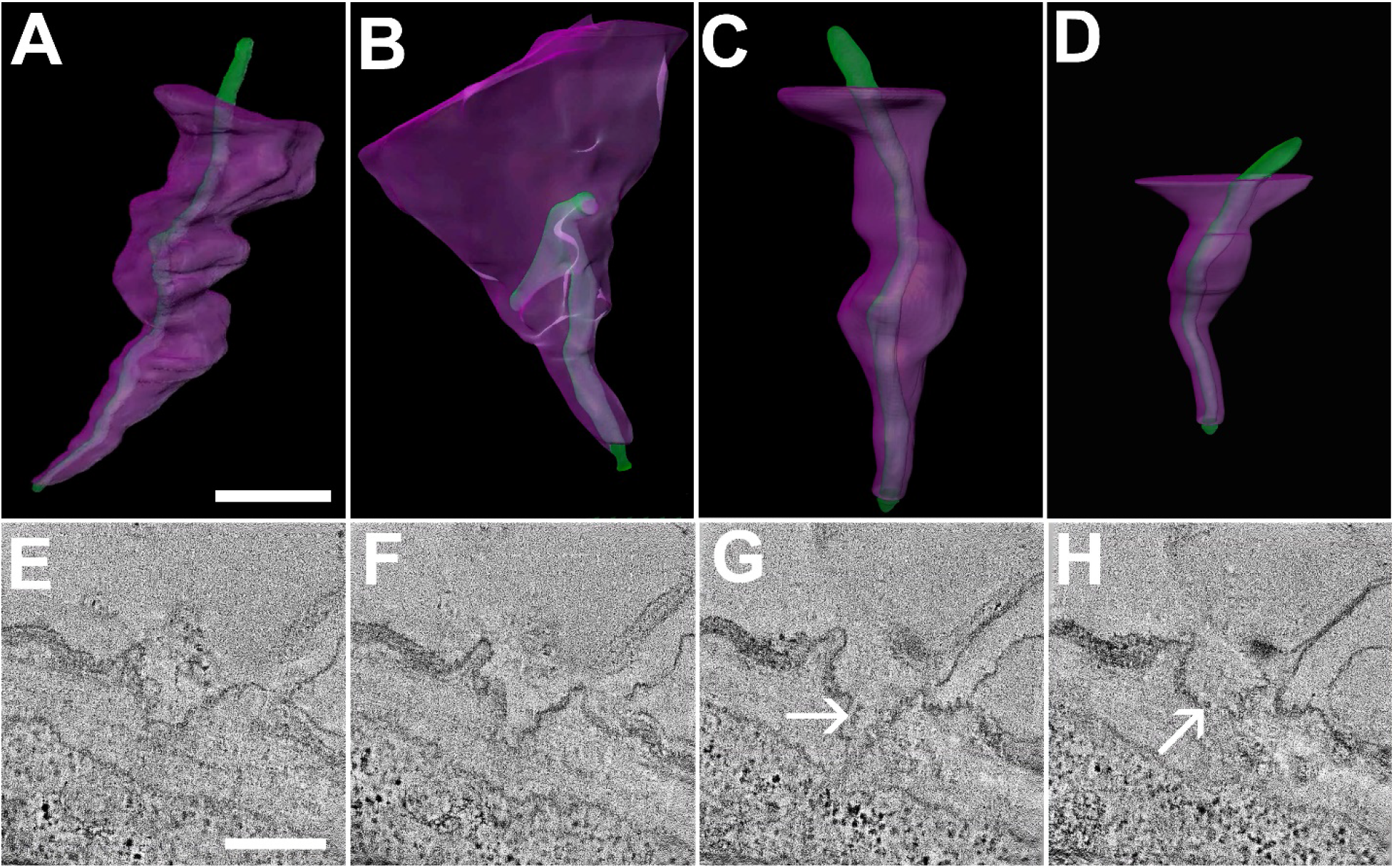
3D reconstructions of funnel plasmodesmata. Electron tomography reconstructions of funnel plasmodesmata from (A) *Triticum aestivum*, (B) *Oryza sativa*, (C) *Medicago sativa*, and (D) *Arabidopsis thaliana*. The lumen of the cell wall pore is rendered in magenta, while the desmotubule appears in green. The shapes of the cell wall pores and their surface structures varied widely in all investigated species. E-H) Four sections of an electron tomograph of a funnel plasmodesma in *Oryza sativa*. The exact dimensions of the desmotubule (arrows) are difficult to determine because the contrast depends on desmotubule location within the sample volume and other parameters. Scale in A = 100 nm (representative for A-D). Scale E = 200 nm (representative for E-H).

In the next step, we used electron tomography surface reconstructions of selected funnel plasmodesmata as templates and extracted tetrahedral grids for theoretical hydrodynamics analyses by finite element methods (Bassi and Rebay 1997). Flow simulations confirmed that flow velocity under a given cell-to-cell gradient of hydrostatic pressure increased with decreasing diameter of the funnel plasmodesmata (Fig. 4). Since the flow rate in a channel equals the pressure gradient divided by hydrodynamic resistance, this implied that overall flow resistance in funnel plasmodesmata is mostly determined by the width of their narrow outlet apertures into the phloem pole pericycle cells.

**Figure 4:**
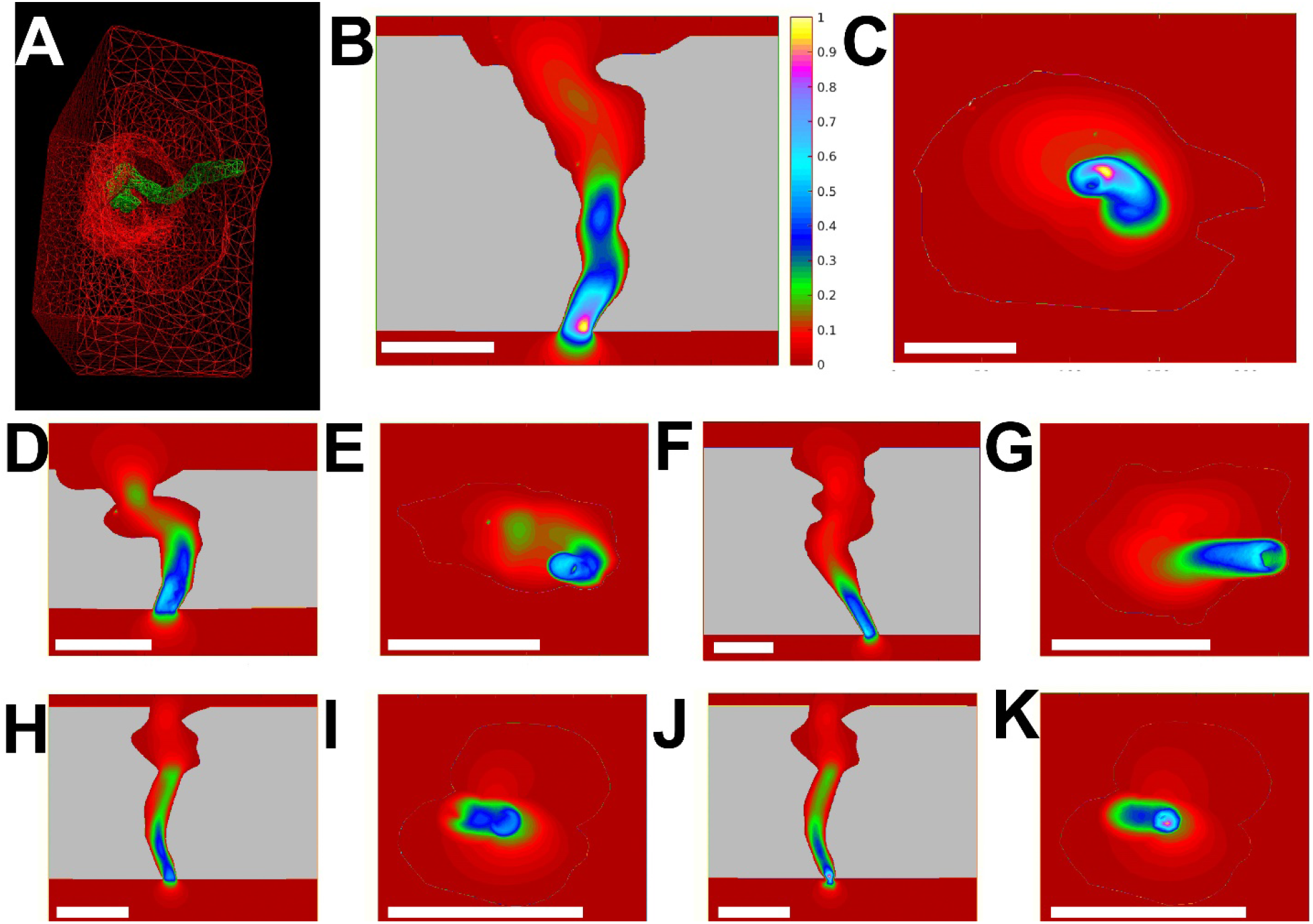
Flow velocity distributions in funnel plasmodesmata determined by fluid dynamics simulations. A) Tetrahedral grit construction of a *Triticum aestivum* funnel plasmodesma (red) including the desmotubule (green), as an example of the input used for the COMSOL software to compute flow patterns in the cytosolic space between plasma membrane and desmotubule. Longitudinal and cross-sectional projections of flow patterns in funnel plasmodesmata of *Ipomoea nil* (B,C), *Oryza sativa* (D,E), and *Triticum aestivum* (F,G). The color scale shows relative values of flow velocity ranging from minimum (red) to maximum (yellow; see scale in B) in each graph. As could be expected, the fastest flow velocities occurred in the narrowest parts of the wall pores, implying that the total hydrodynamic resistance of funnel plasmodesmata depends mostly on the dimensions of the wall pores close to their apertures into the phloem pole pericycle cells. (H-K) Funnel plasmodesma in *Catalpa speciosa*. The electron tomograms did not fully resolve dimensions of the space between plasma membrane and desmotubule. Manual adjustment of the diameter of this cytosol-filled sleeve to 4 nm (H,I) and 8 nm (J,K) yielded almost identical patterns of flow velocity. Scale bar: 100 nm

### Theoretical analysis of funnel geometry effects on diffusion and bulk flow

Since the development of a realistic model describing rates of diffusion and of convective flow as well as electrokinetic effects was beyond the scope of the present study, we restricted further theoretical analysis of the impact of funnel geometries on plasmodesmal transport to the resistances offered by various channel geometries to bulk flow and diffusion. As our standard or control condition, we considered cylindrical channels of radius *a* and length *L* (Fig. 5A, left). A rod of radius *b* in the center of the channels mimicked a desmotubule, and flow and diffusion were assumed to occur only in the annular sleeve between the wall of the channel and the central rod (Fig. 5A, left). In our calculations, we set channel length *L*, equivalent to wall thickness, to 400 nm. While radius *b* was kept constant at 7.5 nm, the outer radius, *a* = *b* + *s*, was varied to obtain sleeve widths (*s*) of 2, 4, and 8 nm (the latter value may seem high, but proteins of 112 kDa, corresponding to a hydrodynamic diameter of ~8 nm, may pass through funnel plasmodesmata; Ross-Elliott et al. 2017). As a result, we had three cylindrical channel models that differed only in radius *a* and thus in sleeve width. These models served as cylindrical standards to which conical channels - i.e., funnel-shaped ones - could be compared. In these conical channels, radius *b* remained constant but radius *a* increased steadily from the smaller, or outlet aperture, toward the larger, or inlet aperture (Fig. 5A, right). Consequently, sleeve width *s* and the cross-sectional sleeve area *A* increased in the same direction as well, in dependence on the magnitude of θ, the angle between the channel wall and the surface of the central rod (Fig. 5A, right). Sleeve widths at the outlet apertures - minimum sleeve widths, in other words - were set to 2, 4, and 8 nm, to allow for direct comparison to the cylindrical channels defined above.

**Figure 5:**
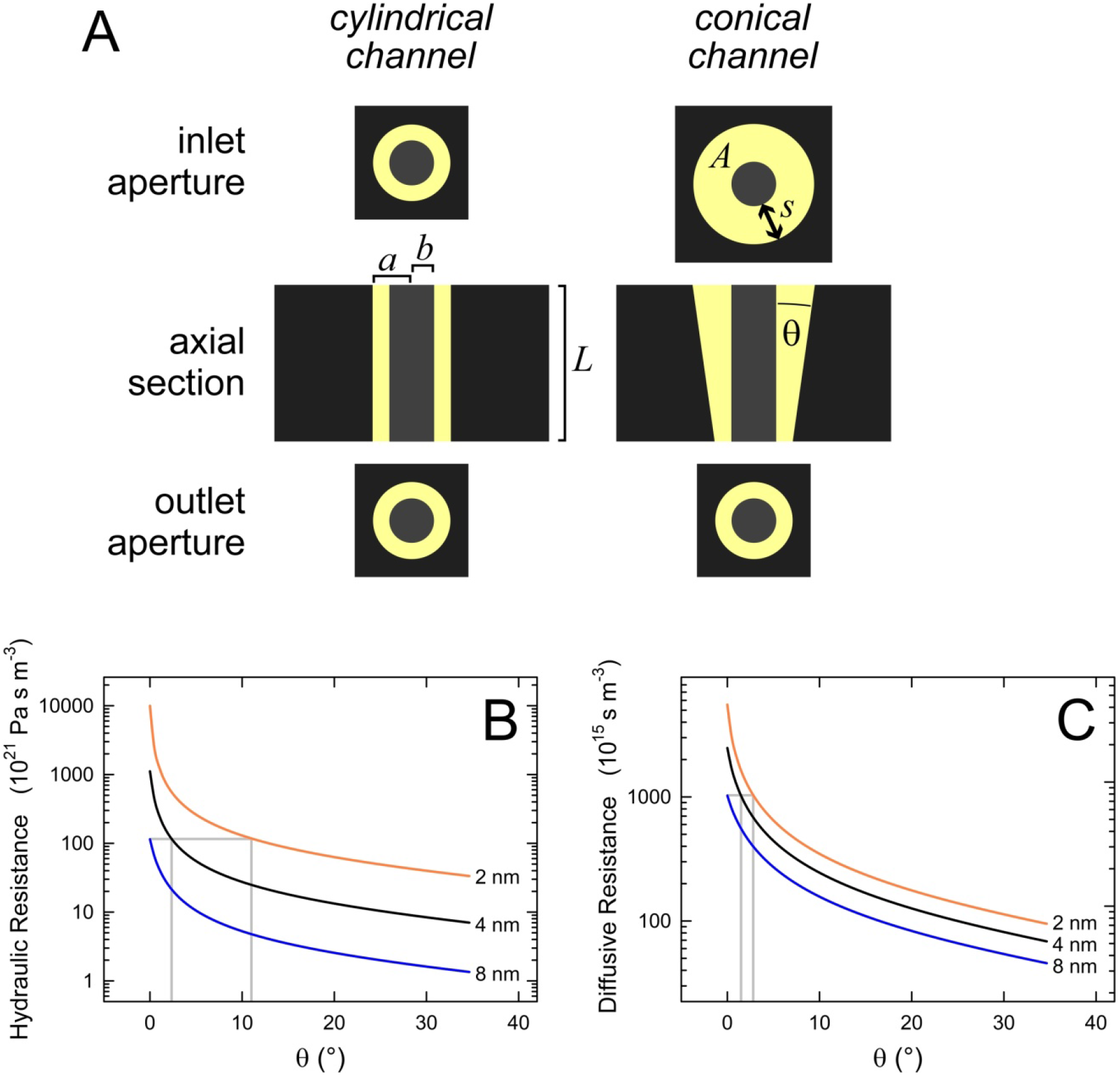
Theoretical evaluation of the effects of channel geometry on hydraulic and diffusive resistance. (A) Models of annular channels with radius *a* in walls of thickness *L*. Transport occurs only in the sleeve (yellow) between the outer channel wall and the central rod of radius *b*. The width of this sleeve (*s*) is constant in cylindrical channels (left) but increases steadily from the outlet to the inlet aperture in conical ones (right), as does the cross-sectional area (*A*) of the sleeve. Results shown below and in Fig. 6 were obtained with *L* (400 nm) and *b* (9.5 nm) kept constant, sleeve width at the outlet aperture (*s*_0_) set to 2 (orange), 4 (black), or 8 nm (blue) by adjusting the pore radius at that position (*a*_0_), and angle θ between channel wall and central rod varying from 0° to 35°. (B) Absolute values of hydraulic resistance as functions of θ. In cylindrical channels (θ = 0°), hydraulic resistance decreases roughly ten-fold with every doubling of the sleeve width. However, channels with minimum sleeve widths (*s*_0_) of 2 and 4 nm reach the resistance of a cylindrical channel with an 8 nm sleeve at relatively small θ, as indicated by grey lines. (C) Diffusive resistances as functions of θ; details as in (B).

The hydraulic resistances of conical channels decreased drastically even with very small angles θ (Fig. 5B). On the other hand, the resistance offered by channels with wider sleeves was substantially lower than that found in narrower ones for all values of θ (Fig. 5B). However, the overlap in the vertical direction of the curves in Fig. 5B indicated that conical geometries in channels with narrow sleeves could reduce the hydraulic resistance to values found in cylindrical channels with much wider sleeves. For instance, the resistance of a cylindrical channel with an 8 nm sleeve was met by conical channels with 4 and 2 nm minimum sleeve widths when θ reached 2.4° and 11°, respectively (highlighted in Fig. 5B). Values of diffusive resistance did not differ as strongly between channels of different sleeve widths for any given angle θ (Fig. 5C) as hydraulic resistance did (Fig. 5B). Presumably, this qualitative difference stems from the fact that diffusive resistance is inversely proportional to the conductive area, whereas hydraulic resistance scales with the inverse area squared (eqs. 8 and 11). As a consequence, conical channels of 4 and 2 nm minimum sleeve width showed the same diffusive resistance as a cylindrical channel with an 8 nm sleeve at angles θ of only 1.5° and 2.8°, respectively (Fig. 5C).

When we normalized the reductions of hydraulic resistance computed for conical channels with respect to the resistances of cylindrical channels of the same sleeve widths, it became clear that the effects were relatively larger in channels with narrower sleeves. For example, hydraulic resistance was reduced to 10% of that found in cylindrical channels of 2, 4, and 8 nm sleeve width in conical channels with angles θ of 1.3°, 2.5°, and 4.6°, respectively, as highlighted in Fig. 6A. The effects of conical geometries on diffusive resistance were qualitatively similar but relatively less pronounced. Reductions of resistance to 10% required angles θ of 5.9°, 9.8°, and 16° in channels of 2, 4, and 8 nm minimum sleeve width, respectively (Fig. 6B). Plots of the ratios of the modulations of hydraulic and diffusive resistances computed for conical model channels (Fig. 7) suggested that relative reductions of hydraulic resistance across the range of cell wall pore opening angles observed in real cells (Fig. 2) were two- to five-fold larger than the corresponding relative changes in diffusive resistance. As a result, convective flow will become more important relative to diffusion when θ increases.

**Figure 6:**
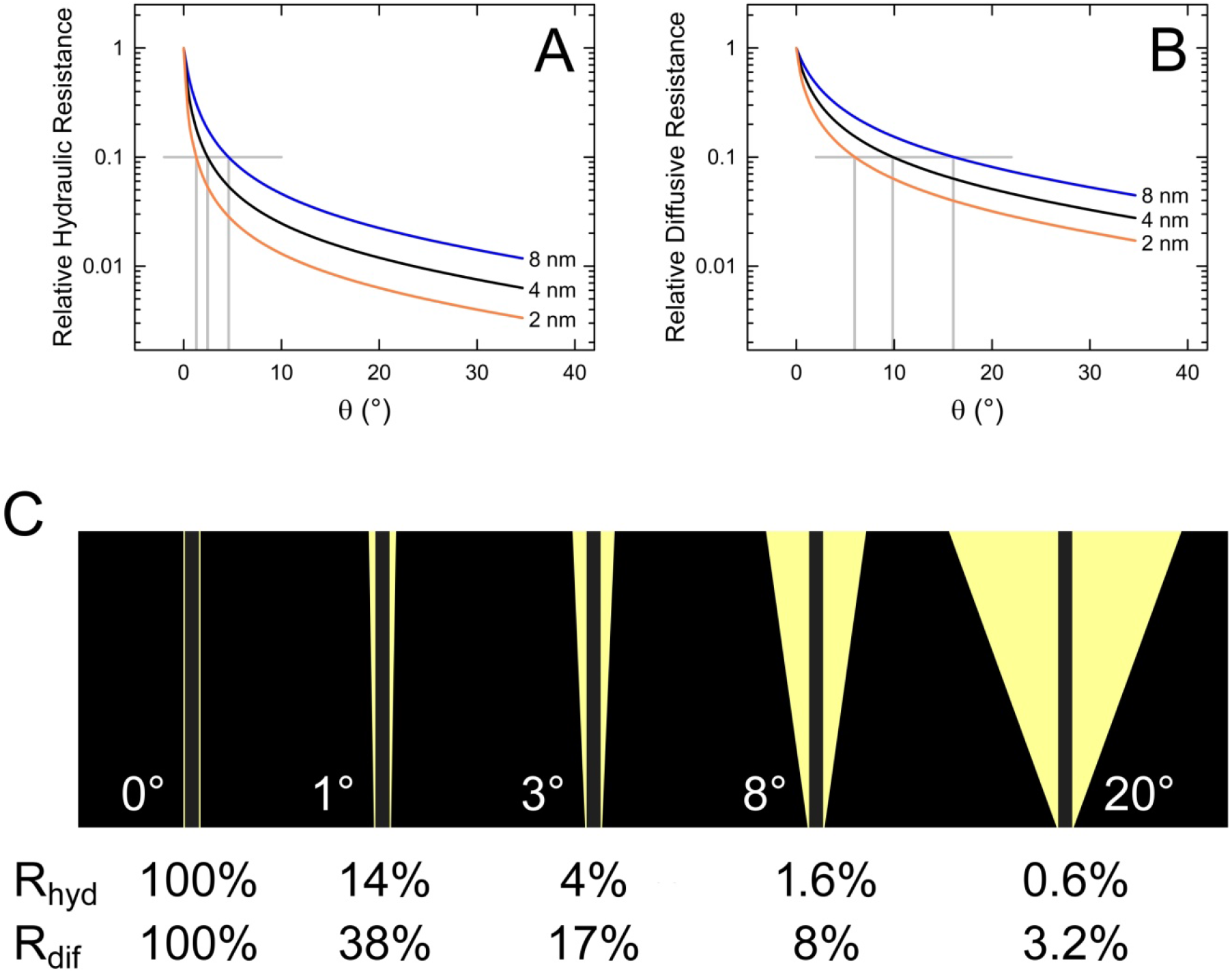
Relative reductions of hydraulic and diffusive resistance in conical channels. (A) Hydraulic resistance of conical channels with varying θ and minimum sleeve widths of 2 (orange), 4 (black), and 8 nm (blue), normalized with respect to the resistance of a cylindrical channel of the corresponding sleeve width. Grey lines highlight angles θ at which the resistance is reduced to 10% of that of conical channels. (B) Dependence of diffusive resistance on θ; details as in (A). (C) Examples of model channels drawn to scale to visualize actual geometries; the minimum sleeve width at the outlet aperture (bottom) is 2 nm, channel length is 400 nm. Values of hydraulic resistance (R_hyd_) and diffusive resistance (R_dif_) are given as percentages of the resistances of the cylindrical control channel (θ = 0°; left).

**Figure 7:**
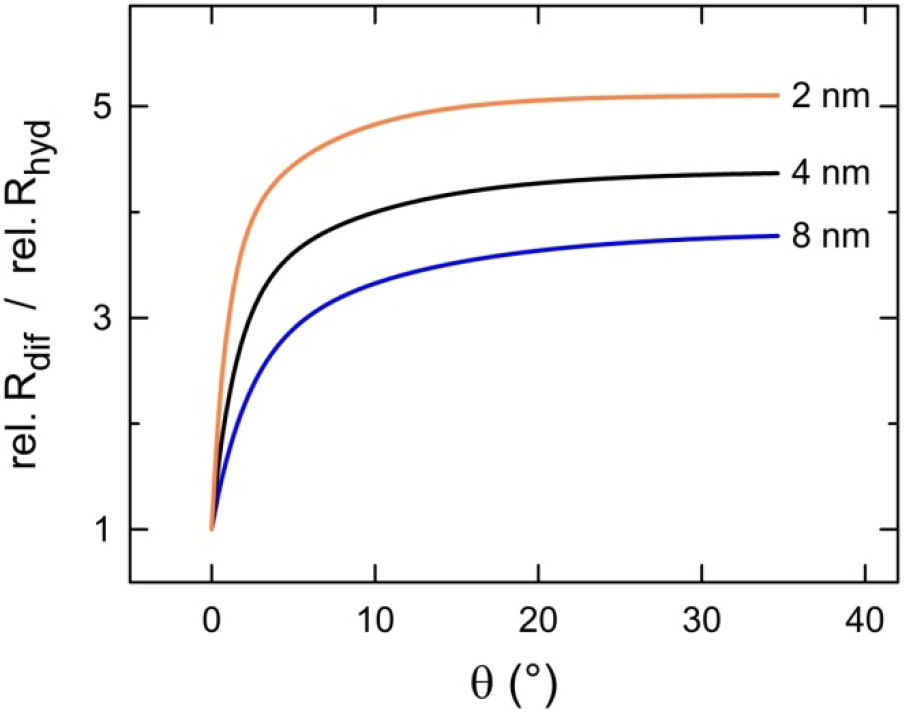
Ratios between the relative change in diffusive resistance (rel. R_dif_; from Fig. 6B) and the relative change in hydraulic resistance (rel. R_hyd_; from Fig. 6A) at different pore angles. With increasing angle θ, the relative hydraulic resistance decreases more strongly than the relative diffusive resistance, implying that convective processes become more important compared to diffusive processes as the angle widens. The effect is stronger in narrower pores.

## Discussion

Most structural and functional plasmodesma studies have been conducted on accessible tissues such as trichomes (Oparka and Prior 1992; Christensen et al. 2009; Barton et al. 2011; Howell et al. 2020), the leaf epidermis (Fitzgibbon et al. 2013), and the calyptra and root meristem (Nicolas et al. 2017). Unfortunately, some of the physiologically most important cases of plasmodesmal transport occur in less accessible tissues, for example the terminal sieve elements of the sieve tubes that deliver the fuel for root growth. These cells are deeply embedded in the central cylinder, which itself is covered by multiple cell layers of the root cortex. We modified fixation techniques for electron microscopy that previously had been applied successfully in *Arabidopsis*, and found funnel plasmodesmata in the phloem unloading zone in root tips of all seven angiosperms examined (Fig. 1).

The geometry of funnel plasmodesmata seemed to differ between species. Estimated opening angles of the cell wall pores clustered around 2°-5° in *C. speciosa* and 15°-22° in *O. sativa*; the other five species showed intermediate values (Fig. 2D). Such structural variability does not necessarily imply significant functional differences. Our theoretical evaluations indicated that small opening angles have comparatively large effects on the hydraulic and diffusive resistance of model channels designed to resemble funnel plasmodesmata structurally (Figs. 5, 6). Compared to cylindrical, simple plasmodesmata, and assuming a width of the cytosolic sleeve of 4 nm, opening angles as found in *C. speciosa* (around 3°) would be expected to reduce the hydraulic resistance by over 90%, and the diffusive resistance by some 70% (Fig. 6A,B). The much larger wall pore opening angles measured in *O. sativa* (around 19°) produce only modestly increased effects, reducing hydraulic resistance by about 98% and diffusive resistance by 94%. Evidently, the physical resistance to symplasmic phloem unloading by bulk flow and diffusion is significantly decreased by the funnel-like shape of some of the plasmodesmata involved in all of the species studied. We emphasize that at standard phloem flow velocities, the entire volume of a sieve element is exchanged within one or a few seconds (Froelich et al. 2011); the rates at which sieve tubes are unloaded obviously have to be commensurate. The specific occurrence of funnel plasmodesmata in the interfaces between protophloem sieve elements and phloem pole pericycle cells supports the idea that these cells provide the main symplasmic route for the required high-capacity phloem unloading.

In this context it seems of interest that for values of θ that correspond to the opening angles estimated for real cells (Fig. 2C), the theoretical reduction in relative hydraulic resistance is two to five times larger than the reduction of diffusive resistance (Fig. 6C). Consequently, the balance between convective and diffusive processes in overall symplasmic transport is expected to shift toward bulk movements when funnel plasmodesmata are formed between cells. This supports the view that rapid phloem unloading in root tips proceeds mainly as bulk flow.

Rather than building structurally complex cytoplasmic bridges such as funnel plasmodesmata, plant cells could form larger numbers of simple plasmodesmata per unit cell wall area or widen the diameters of existing simple plasmodesmata to increase symplasmic transport capacity. We see at least two functional factors that might have favored the evolution of funnel plasmodesmata.

First, many sink organs including root tips are actively growing, which requires complex fine-tuning of the mechanical properties of the expanding cell walls (Cosgrove 2018). Conceivably, increased densities and/or diameters of the pores in expanding cell walls could interfere with the mechanical control of the growth process. Funnel-shaped cell wall pores therefore may represent a compromise between the requirements for rapid symplasmic transport on one hand and the maintenance of the mechanical integrity of the growing cell wall on the other.

Second, as discussed in the Introduction, plasmodesmata function like sieves, preventing particles above a critical size from passing through the narrow cytosolic sleeves. In contrast to cylindrical pores with expanded diameters, funnel plasmodesmata allow for enhanced transport rates while retaining their cargo size-based selectivity. Therefore the presence of funnel plasmodesmata in root tips is in line with the assumption of a physiological necessity for controling the size of particles that leave the sieve tube. Unusually large molecules in the sieve tube stream include ribosomal subunits (Ostendorp et al. 2017) and other cytoplasmic degradation products from developing sieve elements (Knoblauch et al. 2018). The hypothesis that funnel plasmodesmata control the efflux of these and other large molecules from sieve tubes is supported by observations made in *Arabidopsis* roots. There, small molecules moved from sieve elements to phloem pole pericycle cells continuously whereas large proteins entered the latter in a pulsed manner named ‘batch unloading’ (Ross-Elliott et al. 2017). Evidently, the movement of various molecules through funnel plasmodesmata was differentially regulated depending on molecule size. How batch unloading works remains unclear at this time, but thermal motion of the desmotubule could provide a simple but sufficient explanation, according to the hypothetical cargo-gating mechanism (Peters et al. 2021). A tight control of sieve tube efflux based on molecule size might confer advantages to plants battling viruses that utilize the sieve tubes as routes for systemic infection. Generally, virus particles are too large to pass through plasmodesmata, and require specific movement proteins encoded in the viral genome to enter sieve elements (Nelson 2005). In contrast to the mechanisms by which viruses enter sieve tubes, their exit mechanism(s) is mostly unknown (Hipper et al. 2013). Notably, there is no protein synthesizing machinery in sieve elements that a virus could hijack to produce support proteins. By retaining one comparatively narrow aperture, funnel plasmodesmata might at least slow the systemic spread of viruses while massively increasing their conductivity for smaller particles.

Finally, we emphasize that resistance reduction effects similar as described here for funnel-shaped channels must be expected in other partially widened pore structures as well. Plasmodesmata with expanded central cavities, for example, appear quite common in leaf tissues, and can often be found in thickened regions of the cell wall (Robinson-Beers and Evert 1991; Russin and Evert 1985; Oparka et al. 1999; Ehlers and Kollmann 2001). As the example in Fig. 8 and previous studies (Blake 1978; Deinum et al. 2019) demonstrate, plasmodesmata of such an architecture can be more conductive than cylindrical ones of the same minimum sleeve width in much thinner portions of the wall. In funnel plasmodesmata, the very small opening angles that suffice to halve hydraulic and diffusive resistance (Fig. 6A,B) represent structural intricacies that probably will be missed on most TEM micrographs. Even if they were detected, they likely would be neglected in attempts to quantitatively model transport through these pores. Our results suggest that with regard to the efficiency of symplasmic transport, the idea that plasmodesmata can be adequately described as cylindrical or annular tubes might be a seriously misleading simplification. After all, a single funnel-shaped plasmodesma can be as conductive as dozens of cylindrical simple plasmodesmata taken together. In this light it appears necessary to collect precise ultrastructural data concerning plasmodesmata in various cell interfaces to understand and model symplasmic flow through plant tissues. Counting plasmodesma numbers and determining size exclusion limits appears insufficient.

**Figure 8:**
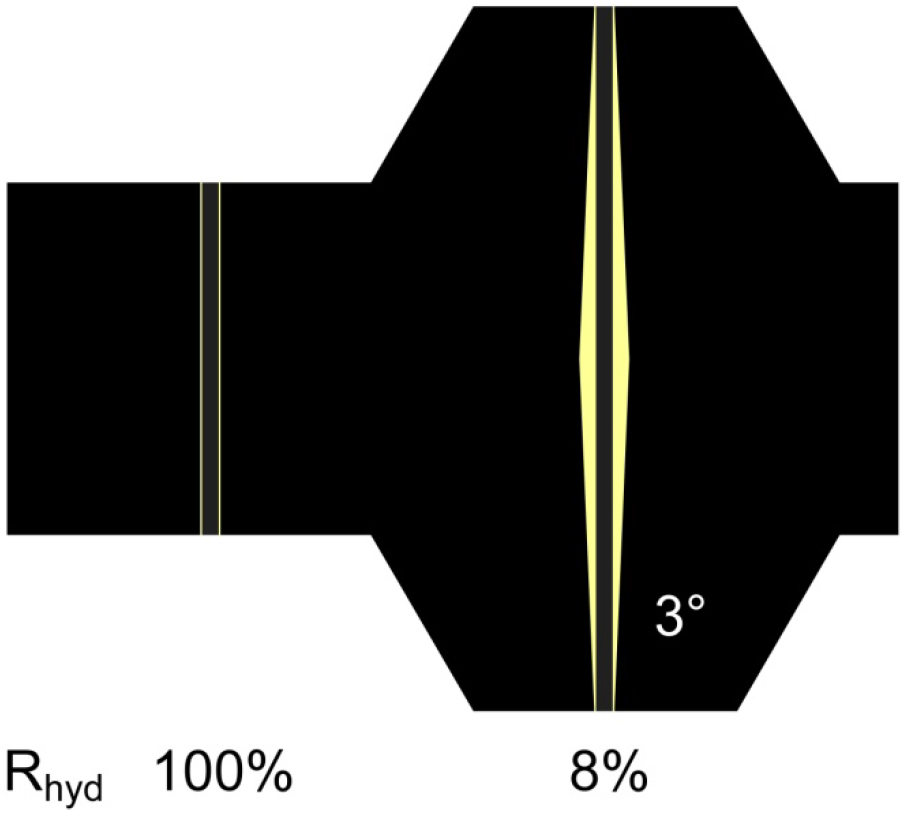
Reduction of hydraulic resistance in a biconical channel. Plasmodesmata with central widenings are often found in thickened portions of the cell wall. This example compares the hydraulic resistance of a biconical model channel (right) with opening angle θ = 3° and apertural sleeve width 2 nm in a thickened cell wall (800 nm) to that of a cylindrical channel (left) with 2 nm sleeve width in a wall of 400 nm thickness. Model channels are drawn to scale to visualize actual geometries.

## Methods

### Plant materials and growth conditions

Two monocots, *Triticum aestivum* and *Oryza sativa* (Poaceae), and the eudicots *Nicotiana tabacum* (Solanaceae), *Ipomoea nil* (Convolvulaceae), *Solenostemon scutellarioides* (Lamiaceae), *Catalpa speciosa* (Bignoniaceae), and *Medicago sativa* (Fabaceae) were grown from seeds in soil, and were maintained in a Greenhouse at 22°C, 60–70% relative humidity, and a 14 h light/10 h dark photoperiod (daylight augmented by 150 μmol m^−2^ s^−1^, Lamp Fixture #PL 90 cv PL Light Systems, Beamsville ON, Canada).

### Sample fixation and embedding

Root tips harvested from mature plants were fixed and embedded based on Wu et al. (2012), but protocols had to be modified for some species. Excised root tips were fixed in 2% paraformaldehyde, 2% glutaraldehyde, 50 mM cacodylate buffer, pH 7 in a microwave oven (Biowave Pro, Pelco, Fresno CA, USA) at 750 W for two 90 s intervals on ice. The samples were washed 3 times for 10 min with distilled water and then post-fixed overnight in 1% OsO_4_. *I. nil*, *S. scutellarioides*, and *M. sativa* were again post-fixed in 2% OsO_4_ for 2 hours, *C. speciosa* for 6 hours, and *T. aestivum* as well as *O. sativa* overnight to receive proper fixation in deeper cell layers. Samples were dehydrated in the microwave oven in a methanol series (up to 80% in 10% steps, 85%, 90%, 95%, 2 × 100% v/v, 1 min each) at 750 W irradiation. The methanol was replaced by propylene oxide in steps (50%, 2 × 100% v/v), and the samples were infiltrated with Spurr’s resin (1:3, 1:2, 1:1, 2:1, 3 × pure resin for 1 day each). Embedded root tips were cured overnight at 60°C. Thin (70 nm) and semi-thin (220–280 nm) sections were produced with a Reichert Ultracut R ultramicrotome (Leica Microsystems, Wetzlar, Germany) and collected on formvar-coated Ni slot grids (Electron Microscopy Sciences, Hatfield PA, USA). Sections were stained with 2% uranyl acetate and 1% potassium permanganate for 12 min, followed by Reynold’s Lead Citrate for 6 min. Semi-thin sections were post-stained with 1% tannic acid. 15 nm colloidal gold feducals were precipitated on both grid surfaces by covering the grids with 5 μL solution for 10 min.

### Transmission electron microscopy and plasmodesma geometry

Micrographs and tomograms were produced using a 200 kV Tecnai G2 20 Twin transmission electron microscope (Thermo Fisher, Waltham MA, USA) equipped with an LaB6 filament and an FEI Eagle 4k CCD camera. Structural parameters including plasmodesma length (*L*; equivalent to cell wall thickness) and the diameters of the plasmodesmal apertures on the sides of the protophloem sieve element and the phloem pole pericycle cell (*D*_PSE_ and *D*_PPP_, respectively) were determined, as far as possible, on electron micrographs using ImageJ/Fiji IJ 1.46r (https://imagej.nih.gov/ij). The opening angle (θ) of the cell wall pore was estimated as

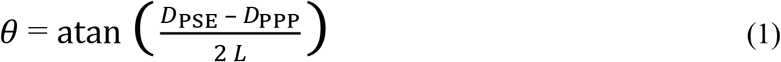

Because *D*_PPP_ could not be determined unambiguously on many micrographs, θ was calculated for *D*_PPP_ = 20 nm as well as *D*_PPP_ = 50 nm, to cover the range of realistic values. Potential correlations between structural parameters were visualized as Geometric Mean Functional Relationships (GMFR; Draper and Smith 1998).

### Electron tomography and 3D modeling

Using the transmission electron microscope described above, tilt series (+55° to −55° in 2° steps) were captured along two single-tilt orthogonal axes with the automated tomography acquisition suite Xplore3D (Thermo Fisher). Raw stacks were combined into dual-axis tomograms (Mastronarde 1997; Kremer et al. 1996) with the open source package IMOD 4.9 (https://bio3d.colorado.edu/imod/). Plasmodesmal tomograms were manually partitioned into two-material image segments comprising the desmotubule and inner pore volume with Amira 6.7 (Thermo Fisher). Tomography surface reconstructions were transferred into tetrahedral grid reconstructions with no more than 18000 triangles and used as input files for the flow simulation software COMSOL Multiphysics 5.4 (COMSOL Multiphysics, Burlington MA, USA).

### Theory and computational analysis

We used a combination of numerical simulations and theory to evaluate the transport properties of funnel plasmodesmata. The analysis of diffusion was based on Fick’s law, which links the flux, *j*, the concentration gradient ∇*c*, and the diffusion constant *D* (which for modeling purposes was assumed to be 6.7 × 10^−10^ m^2^s^−1^):

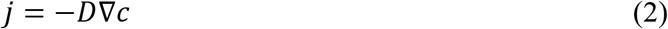

Assuming solute conservation and steady-state conditions leads to the diffusion equation

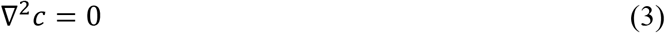

Transport characteristics of plasmodesmatal geometries extracted from Amira 6.7 were modeled using COMSOL Multiphysics 5.4. The diffusion current *I* was determined by integrating the flux across the pore entrance. The diffusion resistance

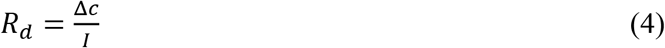

was computed where Δ*c* is the imposed cell-to-cell concentration difference. In an idealized concentric geometry, the current *I* can be expressed as

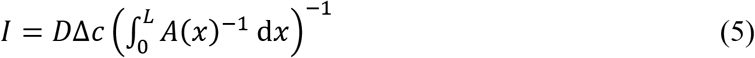

where *L* is the length of the pore and *A*(x) is the cross-sectional area of the open space in the pore, measured as a function of the distance along the pore axis, *x*. For models with a central rod of constant diameter *b* mimicking the desmotubule as shown in Fig. 5A, the outer radius *a* also is a function of *x*:

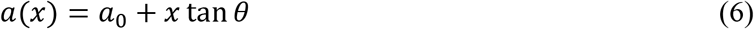

where *a*_0_ refers to the smaller, or outlet aperture and *θ* is the pore angle (see Fig. 5A).

Consequently, the cross-sectional area *A* of the space available for transport, or sleeve, changes along the pore axis *x* according to

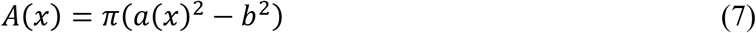

Combining Eqs. (4) and (5) leads to an expression for the diffusion resistance:

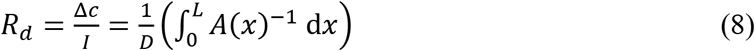

The analysis of bulk flow through the pores was based on the Stokes equation,

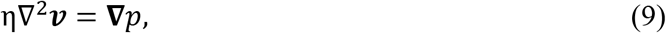

where *v* is the velocity field, *p* is pressure, and η is the cytoplasmic viscosity (for modeling purposes, η was set to 8.9 × 10^−4^ Pa s). A pressure drop Δ*p* was applied across the pore and we assumed no-slip conditions (*v* = 0) on all solid boundaries.

Flow characteristics of plasmodesmatal geometries extracted from Amira 6.7 were modeled using COMSOL Multiphysics 5.4. Validation of the solver was carried out as described by Jensen et al. (2012). The volumetric flow rate *Q* was determined by integrating the velocity field across the pore entrance. Subsequently, the hydraulic resistance

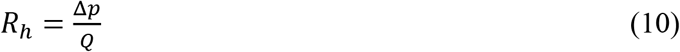

was computed. In an idealized concentric, conical geometry, the hydraulic resistance can be expressed as

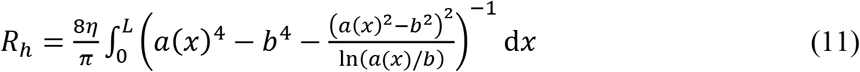

Both the numerical solutions of the COMSOL flow/diffusion analyses and physical models comparing plasmodesmatal structural elements to theoretical flow profiles were examined using MATLAB R2020b (MathWorks, Natick MA, USA).

## References

Barton DA, Cole L, Collings DA, Liu DYT, Smith PMC, Day DA, Overall RL. 2011. Cell-to-cell transport via the lumen of the endoplasmic reticulum. Plant Journal 66:806–817. DOI10.1111/j.1365-313X.2011.04545.x

Bassi F, Rebay S. 1997. A high-order accurate discontinuous finite element method for the numerical solution of the compressible Navier-Stokes equations. Journal of Computational Physics 131:267–279. DOI:10.1006/jcph.1996.5572

Blake JR. 1978. On the hydrodynamics of plasmodesmata. Journal of Theoretical Biology 74:33–47. DOI:10.1016/0022-5193(78)90288-6

Christensen NM, Faulkner C, Oparka K. 2009. Evidence for unidirectional flow through plasmodesmata. Plant Physiology 150:96–104. DOI:10.1104/pp.109.137083

Comtet J, Turgeon R, Stroock AD. 2017. Phloem loading through plasmodesmata: a biophysical analysis. Plant Physiology 175:904–915. DOI:10.1104/pp.16.01041

Cosgrove DJ. 2018. Diffusive growth of plant cell walls. Plant Physiology 176:16–27. DOI:10.1104/pp.17.01541

Deinum EE, Mulder BM, Benitez-Alfonso Y. 2019. From plasmodesma geometry to effective symplasmic permeability through biophysical modelling. eLife 8:e49000. DOI:10.7554/eLife.49000

Ding B, Turgeon R, Parthasarathy MV. 1992. Substructure of freeze-substituted plasmodesmata. Protoplasma 169:28–41. DOI: 10.1007/Bf01343367

Draper NR, Smith H. 1998. Applied Regression Analysis, 3^rd^ ed. John Wiley and Sons, New York.

Ehlers K, Kollmann R. 2001. Primary and secondary plasmodesmata: structure, origin, and functioning. Protoplasma 216:1–30. DOI:10.1007/Bf02680127

Esau K, Thorsch J. 1985. Sieve Plate Pore and Plasmodesmata, the Communications Channels of the Symplast: Ultrastructural Aspects and Developmental Relations. American Journal of Botany 72:1641–1653. DOI:10.1002/j.1537-2197.1985.tb08429.x

Fitzgibbon J, Beck M, Zhou J, Faulkner C, Robatzek S, Oparka K. 2013. A developmental framework for complex plasmodesmata formation revealed by large-scale imaging of the *Arabidopsis* leaf epidermis. Plant Cell 25:57–70. DOI:10.1007/Bf01281999

Fitzgibbon J, Bell K, King E, Oparka K. 2010. Super-resolution imaging of plasmodesmata using three-dimensional structured illumination microscopy. Plant Physiology 153:1453–1463. DOI:10.1104/pp.110.157941

Froelich DR, Mullendore DL, Jensen KH, Ross-Elliott TJ, Anstead JA, Thompson GA, Pélissier HC, Knoblauch M. 2011. Phloem ultrastructure and pressure flow: sieve-element-occlusion-related agglomerations do not affect translocation. Plant Cell 23:4428–4445. DOI:10.1105/tpc.111.093179

Gerlitz N, Gerum R, Sauer N, Stadler R. 2018. Photoinducible DRONPA-s: a new tool for investigating cell-cell connectivity. Plant Journal 94:751–766. DOI:10.1111/tpj.13918

Goodwin PB, Shepherd V, Erwee MG. 1990. Compartmentation of fluorescent tracers injected into the epidermal cells of *Egeria densa* leaves. Planta 181:129–136 DOI:10.1007/BF00202335

Hipper C, Brault V, Ziegler-Graff V, Revers F. 2013. Viral and cellular factors involved in phloem transport of plant viruses. Frontiers in Plant Science 4:154. DOI:10.3389/fpls.2013.00154

Howell AH, Peters WS, Knoblauch M. 2020. The diffusive injection micropipette (DIMP). Journal of Plant Physiology 244:153060. DOI:10.1016/j.jplph.2019.153060

Jensen KH, Mullendore DL, Holbrook NM, Bohr T, Knoblauch M, Bruus H. 2012. Modeling the hydrodynamics of phloem sieve plates. Frontiers in Plant Science 3:151. DOI:/10.3389/fpls.2012.00151

Knoblauch M, Peters WS, Bell K, Ross-Elliott TJ, Oparka KJ. 2018. Sieve-element differentiation and phloem sap contamination. Current Opinion in Plant Biology 43:43–49. DOI:10.1016/j.pbi.2017.12.008

Kremer JR, Mastronarde DN, McIntosh JR. 1996. Computer visualization of three-dimensional image data using IMOD. Journal of Structural Biology 116:71–76. DOI:10.1006/jsbi.1996.0013

Lee JY, Frank M. 2018. Plasmodesmata in phloem: different gateways for different cargoes. Current Opinion in Plant Biology 43:119–124. DOI:10.1016/j.pbi.2018.04.014

Liesche J, Schulz A. 2012. Quantification of plant cell coupling with three-dimensional photoactivation microscopy. Journal of Microscopy 247:2–9. DOI:10.1111/j.1365-2818.2011.03584.x

Liesche J, Schulz A. 2013. Modeling the parameters for plasmodesmal sugar filtering in active symplasmic phloem loaders. Frontiers in Plant Science 4:207. DOI:10.3389/fpls.2013.00207

Mastronarde DN. 1997. Dual-axis tomography: an approach with alignment methods that preserve resolution. Journal of Structural Biology 120:343–352. DOI: https://doi.org/10.1006/jsbi.1997.3919

Nelson RS. 2005. Movement of viruses to and through plasmodesmata. In: Oparka K, editor. Plasmodesmata. Dundee: Blackwell Publishing. pp. 188–211. DOI:10.1002/9780470988572.ch9

Nicolas WJ, Grison MS, Trépout S, Gaston A, Fouché M, Cordelières FP, Oparka K, Tilsner J, Brocard L, Bayer EM. 2017. Architecture and permeability of post-cytokinesis plasmodesmata lacking cytoplasmic sleeves. Nature Plants 3:17082. DOI: 10.1038/nplants.2017.82

Oparka KJ, Prior DAM. 1992. Direct evidence for pressure-generated closure of plasmodesmata. Plant Journal 2:741–750. DOI:10.1111/j.1365-313X.1992.tb00143.x

Oparka KJ, Roberts AG, Boevink P, Cruz SS, Roberts I, Pradel KS, Imlau A, Kotlizky G, Sauer N, Epel B. 1999. Simple, but not branched, plasmodesmata allow the nonspecific trafficking of proteins in developing tobacco leaves. Cell 97:743–754. DOI:10.1016/S0092-8674(00)80786-2

Oparka KJ, Turgeon R. 1999. Sieve elements and companion cells–traffic control centers of the phloem. Plant Cell 11:739–750. DOI:10.1105/tpc.11.4.739

Ostendorp A, Pahlow S, Krüßel L, Hanhart P, Garbe MY, Deke J, Giavalisco P, Kehr J. 2017. Functional analysis of *Brassica napus* phloem protein and ribonucleoprotein complexes. New Phytologist 214:1188–1197. DOI:10.1111/nph.14405

Park K, Knoblauch J, Oparka K, Jensen KH. 2019. Controlling intercellular flow through mechanosensitive plasmodesmata nanopores. Nature Communications 10:3564. DOI:10.1038/s41467-019-11201-0

Peters WS, Jensen KH, Stone HA, Knoblauch M. 2021. Plasmodesmata and the problems with size: interpreting the confusion. Journal of Plant Physiology 257:153341. DOI:10.1016/j.jplph.2020.153341

Robinson-Beers K, Evert RF. 1991. Fine structure of plasmodesmata in mature leaves of sugarcane. Planta 184:307–318. DOI:10.1007/BF00195331

Ross-Elliott TJ, Jensen KH, Haaning KS, Wager BM, Knoblauch J, Howell AH, Mullendore DL, Monteith AG, Paultre D, Yan D, Otero S, Bourdon M, Sager R, Lee JY, Helariutta Y, Knoblauch M, Oparka KJ. 2017. Phloem unloading in Arabidopsis roots is convective and regulated by the phloem-pole pericycle. eLife 6:e24125. DOI:10.7554/eLife.24125

Russin WA, Evert RF. (1985) Studies on the leaf of *Populus deltoides* (Salicaceae): ultrastructure, plasmodesmatal frequency, and solute concentrations. American Journal of Botany 72:1232–1247. DOI:10.1002/j.1537-2197.1985.tb08377.x

Rutschow HL, Baskin TI, Kramer EM. 2011. Regulation of solute flux through plasmodesmata in the root meristem. Plant Physiology 155:1817–1826. DOI:10.1104/pp.110.168187

Schulz A. 1999. Physiological control of plasmodesmal gating. In: van Bel AJE, van Kesteren WJP, editors. Plasmodesmata. Berlin, Heidelberg: Springer. pp. 173–204. DOI: 10.1007/978-3-642-60035-7_11

Waigmann E, Turner A, Peart J, Roberts K, Zambryski P. 1997. Ultrastructural analysis of leaf trichome plasmodesmata reveals major differences from mesophyll plasmodesmata. Planta 203:75–84. DOI:10.1007/s004250050167

Wang X, Luna GR, Arighi CN, Lee JY. 2020. An evolutionarily conserved motif is required for Plasmodesmata-located protein 5 to regulate cell-to-cell movement. Communications Biology 3:291. DOI:10.1038/s42003-020-1007-0

Warmbrodt RD. 1985. Studies on the root of *Hordeum vulgare* L. – ultrastructure of the seminal root with special reference to the phloem. American Journal of Botany 72:414–432. DOI:10.2307/2443534

Wu S, Baskin TI, Gallagher KL. 2012. Mechanical fixation techniques for processing and orienting delicate samples, such as the root of *Arabidopsis thaliana*, for light or electron microscopy. Nature Protocols 7:1113–1124. DOI:10.1038/nprot.2012.056

